# Developmental and transcriptional programs that define the initiation of the forelimb

**DOI:** 10.1101/2025.11.23.689776

**Authors:** Vighnesh Ghatpande, Alecxander J. Lewis, Kathryn E. Windsor, Hyunji Lee, Aaron J. Alcala, Douglas B. Menke, Can Cenik, Steven A. Vokes

## Abstract

The vertebrate forelimb initiates as a localized swelling in the somatic lateral plate mesoderm (somatic LPM) in response to TBX5-dependent transcription. The molecular pathways driving limb morphogenesis have been extensively studied but the steps directly preceding limb bud formation remain poorly characterized. To address this, we defined the temporal onset of forelimb initiation in mouse embryos using sequencing based high-throughput approaches (RNA-seq, scRNA-seq, ChIP-seq, and Ribo-ITP) benchmarked to known features in forelimb development, identifying four distinct stages. Using scRNA-seq at the onset of forelimb-specific transcription, we determined the transcriptional profile of the somatic LPM and identified signature genes that distinguish the nascent forelimb from other cell types. This group includes multiple genes involved in neural projection as well as cell adhesion. Interestingly, these genes are highly enriched for TBX5 binding sites, suggesting they are candidate early transcriptional targets of TBX5. As TBX5 is essential for forelimb outgrowth, the identification of these genes suggests new mechanistic models for TBX5-driven limb initiation.

## INTRODUCTION

The vertebrate limb buds, the embryonic structures that give rise to the arms and legs, emerge as swellings from the somatic lateral plate mesoderm (LPM). Initially consisting of mesoderm encased by ectoderm, they undergo substantial growth and morphogenesis before the later migration of angioblasts, myoblasts and neural cells. The limb bud has long been a model system for understanding early inductive signaling processes and has been characterized on morphological and genetic levels. These studies have defined the major genes and pathways that initiate limb development as well as some of the major morphological processes. However, the precise timing and sequence by which these genes initiate limb development remain unclear.

Several staging systems indicate that the mouse forelimb bud, which forms temporally before the hindlimb, is first visible around E9.5 (Fernández-Terán et al., 2006; Martin, 1990; Musy et al., 2018; Wanek et al., 1989). These studies focus on stages occurring during and after the identification of a visible limb bud. However, visible limb bud morphogenesis is preceded by earlier events that were not described in these studies. For example, the somatic LPM undergoes an epithelial to mesenchymal transformation (EMT), during which they lose epithelial characteristics that were established post-gastrulation (Gros and Tabin, 2014). This change in adhesive properties presumably results in increased cell mobility that enables limb bud outgrowth. In zebrafish embryos, somatic LPM cells coalesce to form the pectoral fin through directional migration (Ahn et al., 2002; Mao et al., 2015). LPM cells have also been observed to stream into early mouse limb buds (Wyngaarden et al., 2010), suggesting the possibility of a similar mechanism in mice.

The molecular trigger that initiates EMT in the somatic LPM remains unknown. This process could involve initiating the expression of genes that transcriptionally initiate EMT such as those driving neural crest initiation. In line with this, PRRX1 and TWIST have been proposed to initiate somatic LPM EMT in chick embryos (Newton and Smith, 2024; Newton et al., 2022). Alternatively, translational regulation is a key regulator of EMT in several types of cancer. For example, TGF-beta triggers the translation SNAIL, which is necessary for inducing EMT in mouse models of lung metastasis (Ye et al., 2015). SNAIL is a core transcriptional factor that promotes EMT in the context of cancer and mesoderm migration but is dispensable for neural crest cell (NCC) migration (Carver et al., 2001; Murray and Gridley, 2006). Additionally, RNA binding proteins YB-1 and CELF1 regulate the translation of EMT-inducing factors by promoting cap-independent initiation and by binding to the 3’UTR, respectively (Chaudhury et al., 2016; Evdokimova et al., 2009). Finally, translational elongation is regulated in several EMT contexts. For example, SNAIL, DAB*2,* and the EMT markers VIM and RAC1 are all induced by enhanced translation (Ebright et al., 2020; Hussey et al., 2011; Wurth et al., 2016). Despite the established roles of translational regulation in EMT across systems, to what extent such mechanisms contribute to limb bud EMT remains unknown.

*Tbx5*, the earliest known forelimb-specific marker, is essential for forelimb outgrowth and the causative gene underlying Holt Oram syndrome (Vanlerberghe et al., 2019). It directly activates *Fgf10*, which is also essential for limb bud outgrowth (Agarwal et al., 2003; Delgado et al., 2021; Lizarraga et al., 1999; Min et al., 1998; Nishimoto et al., 2015; Rallis et al., 2003; Sekine et al., 1999; Xu et al., 1998). Despite being broadly expressed at later stages, *Tbx5* is only required prior to or at limb bud initiation, indicating a transient inductive role (Hasson et al., 2007). This therefore defines a critical time window for *Tbx5* during the very earliest stages of limb specification. Both *Tbx5* and its target *Fgf10* have been suggested to promote EMT since in embryos lacking each either of these genes the mesenchyme in the forelimb field is reduced and retains markers of polarized mesenchymal cells (Gros and Tabin, 2014). *Tbx5* paralogues in Zebrafish regulate the directional migration of LPM cells that migrate directionally to form the nascent limb bud (Boyle-Anderson et al., 2022; Mao et al., 2015).

Collectively, these studies indicate a critical role for TBX5 in forelimb development. While the primary role of TBX5 in limb development is thought to be activating *Fgf10*, it is unclear if it acts upstream of other developmental regulators as well. Some studies have observed that unlike *Tbx5^-/-^* embryos, *Fgf10^-/-^* embryos still initiate limb buds, suggesting additional roles for TBX5 in forelimb initiation (Hasson et al., 2007; Sekine et al., 1999). Such roles might include the activation of genes or processes involved in EMT or cell migration that enable forelimb formation. Since this period remains poorly characterized, we first defined a series of developmental stages in forelimb specification that are benchmarked to somite numbers. We then identified signature genes in the stage comprising the nascent limb field and found that most of these harbor TBX5 binding regions, suggesting that they are amongst the earliest direct TBX5 target genes. This gene set is enriched for cell adhesion components, suggesting possible mechanisms by which TBX5 enables forelimb formation.

## RESULTS

### Temporal onset of the expression of limb specific genes

Given its role as the most upstream known forelimb specifying gene, we first examined expression of *Tbx5*, focusing on the LPM adjacent to somites 7-13, which corresponds to the axial level of the future forelimb (Wanek et al., 1989). At 10 somites, *Tbx5* was faintly detected in the LPM. Expression levels were slightly higher at 12 and increased sharply between 14-16 somites (Fig. 1A-D). As TBX5 is thought to induce the expression of Fgf10 (Agarwal et al., 2003; Sekine et al., 1999), we examined *Fgf10*, expecting to see initiation of expression around 12 somites. Unexpectedly, *Fgf10* was present in all stages we examined, starting with 7 somites, where it was expressed throughout the entirety of the LPM (Fig. 1E, n=3). The expression of *Fgf10* was maintained in the LPM at the axial level of the forelimb at subsequent stages (Fig. 1F-L), with upregulated expression at 16S (Fig.1K). This expression pattern is consistent with expression reported in mouse and chick embryos at this stage (Ohuchi et al., 1997; Urness et al., 2011) and suggests that TBX5 does not initiate *Fgf10* expression but rather acts to maintain it within the nascent forelimb. This early expression of *Fgf10* was likely overlooked in prior limb studies, which focused on embryos at stages after limb bud initiation would already have begun (Agarwal et al., 2003; Nishimoto et al., 2015; Rallis et al., 2003). Despite this early expression, *Fgf10^-/-^*embryos have *Tbx5* expression domains that are indistinguishable from control littermates at 12 somites and 16 somites (Fig. S1).

**Figure 1:**
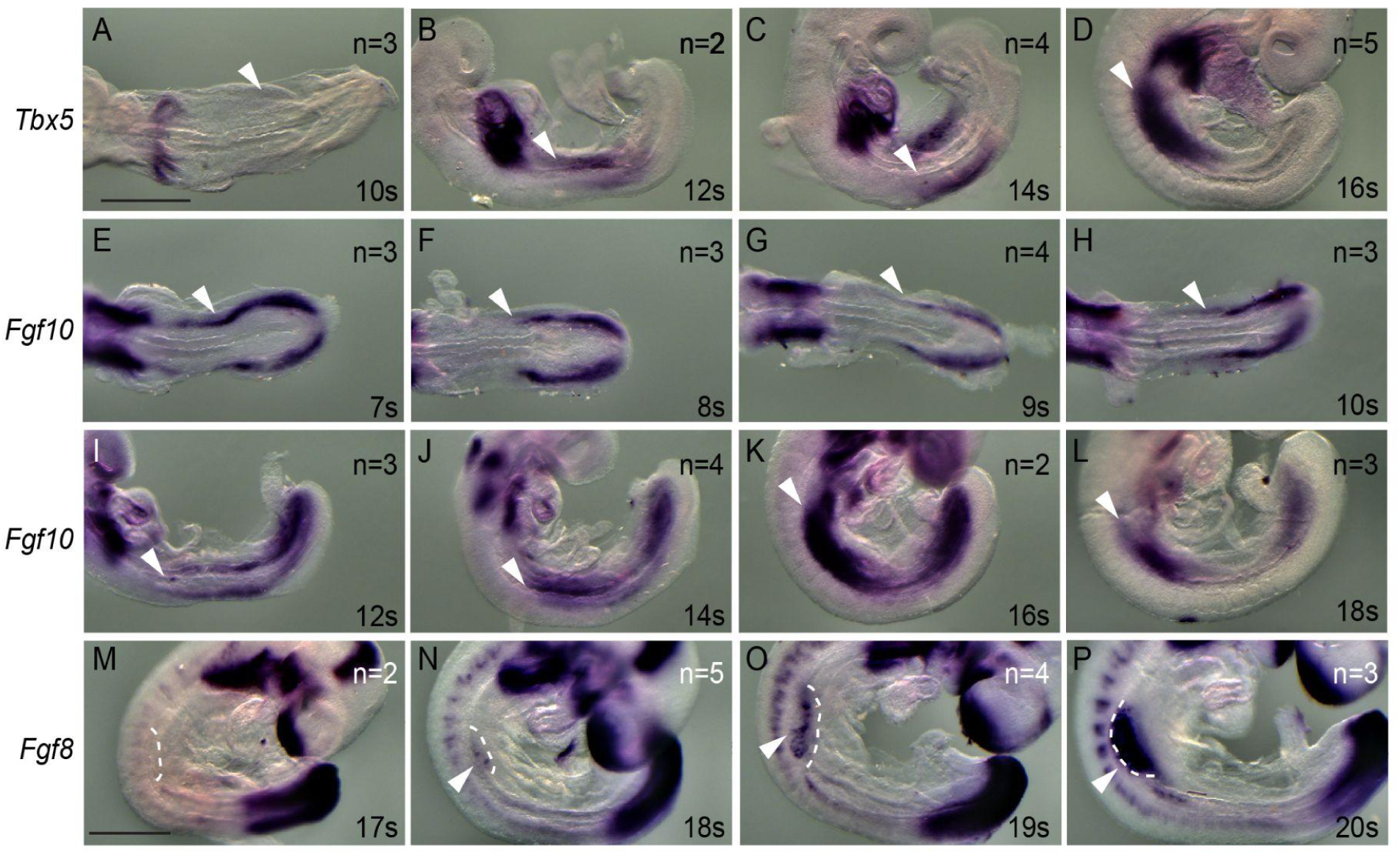
Temporal onset of limb specifier genes. (A-P) In situ hybridization showing expression patterns for the genes listed to the left of each column. White arrows indicate the position of the seventh somite, corresponding to the rostral boundary of the presumptive forelimb field. The embryonic stage in somites #s (s) is indicated at the bottom right of each image and the contour of the forelimb is shown in dashed white lines in (M-P). n refers to the number of biological replicates; the scale bar indicates 500μm.

Having established that *Tbx5* and *Fgf10* are both upregulated at 16 somites, we sought to define the onset of ectodermal *Fgf8* expression. *Fgf8* is genetically downstream of Fgf10 (Barrow et al., 2003; Kawakami et al., 2001; Soshnikova et al., 2003) and it initiates reciprocal signaling between the ectoderm and mesenchyme that signifies a later stage of limb bud formation, ultimately leading to the acquisition of distal polarity. Consistent with prior reports, *Fgf8* is first detectable in ventral ectoderm (Bell et al., 1998; Loomis et al., 1998) in some 18 somite embryos (3/10 forelimbs, n = 5 embryos, Fig. 1N) and is present in the ectoderm of all embryos by 19 somites (6/6 forelimbs, n = 3 embryos, Fig. 1P). Based on this expression we conclude that forelimb-specific gene expression in limb buds initiates between 14-16 somites, preceding the activation of ectodermal *Fgf8* at 18 somites.

### Somatic LPM undergoes a loss of cell polarity between 12-13 somites

The somatic LPM re-epithelializes shortly after gastrulation and then undergoes an EMT in which cells lose markers of cell polarity and subsequently are found within the limb bud. These movements, which are necessary for limb bud morphogenesis, are at least partially dependent on *Fgf10* and *Tbx5* (Gros and Tabin, 2014). To determine the initial onset of this process, we first examined the expression of TWIST, which is expressed early in the somatic LPM in chick, where it is proposed to act as a trigger of EMT (Newton et al., 2022). TWIST (the antibody detects TWIST1 and likely cross-reacts with TWIST2) was broadly expressed in the somatic LPM at 7-8 somites when most of the somatic LPM consists of a single, pseudostratified layer of cells and continued to be expressed at 10 somites when multiple cell layers are present (Fig. S2A-D). We focused our subsequent analysis starting at 10 somites, when *Tbx5* has not yet been upregulated (Fig. 1A).

The somatic LPM of 10 somite embryos uniformly exhibited apical localization of N-Cadherin in the medial layer of cells lining the coelom while it was absent from the more lateral layers of cells (4/4 embryos, Fig. 2A,D). In contrast, most embryos no longer had polarized N-Cadherin at 13 or 15 somites (3/4 and 2/3 embryos, respectively did not contain polarized expression; Fig. 2B,C). Similarly, the somatic LPM contained polarized β-catenin at 10-11 somites (4/6 embryos) while expression was reduced or absent at 13 and 15 somites (3/4 and 3/3 embryos, respectively; Fig. 2G-I). In contrast, aPKC was not polarized in the somatic LPM at 10-11 somites (0/7 embryos) or at 13 and 15 somites (0/4 and 0/4 embryos, respectively; Fig. 2J-L). As the lack of polarized aPKC conflicts with a prior reports, especially in chick (Gros and Tabin, 2014; Newton et al., 2022), we used multiplex imaging to establish that sections contained *Tbx5*, confirming that they are on the axial level of the forelimb field (Fig. 2M-O). The same embryos displayed robust expression of polarized aPKC the epithelialized somite, confirming the effectiveness of the antibody under our conditions (Fig. S2E-H). Interestingly, LAMININ was present within the somatic LPM at all stages examined, although this expression was variable (4/8, 3/4 and 4/4 embryos at 10-11, 13 and 15 somites, respectively; Fig. 2P-R). However, under our experimental conditions, it was not possible to resolve a basal layer of Laminin from the somatic LPM from the basal ectodermal layer and to define a change in LAMININ expression. Even though broader set of junctional markers remain to be assessed, loss of apical N-Cadherin and Beta-catenin is consistent with the conclusion of EMT-like process between 12-13 somites.

**Figure 2:**
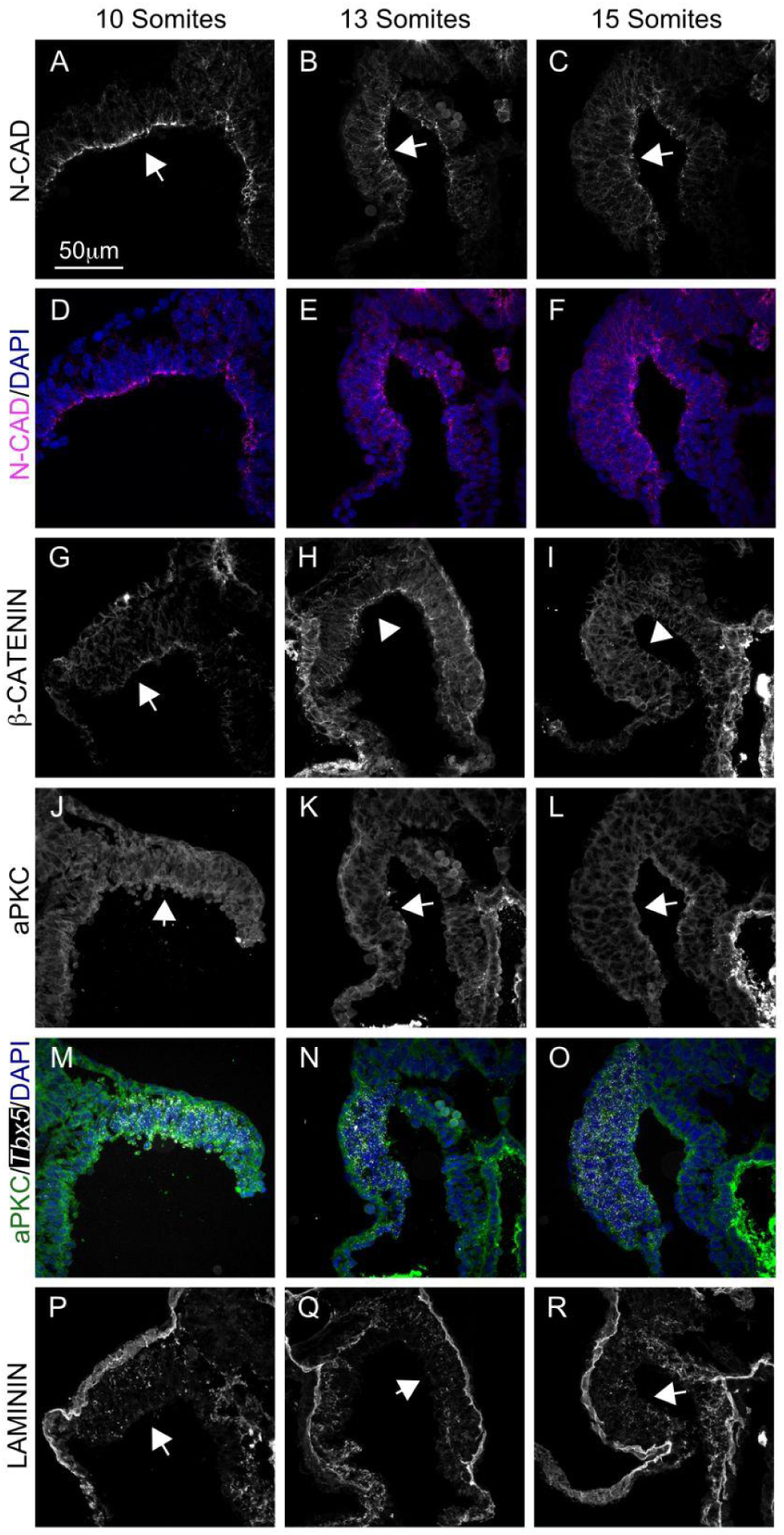
Changes in somatic LPM cell polarity occur between 10-15 somites. (**A-R**) Cryosections showing expression of the indicated markers at 10 somites (left column), 13 somites (middle column) and 15 somites (right column). Arrows point to the apical side of the somatic LPM. (**A-F**) N-Cadherin expression. (**G-I**) β-Catenin expression. (**J-O**) Dual immunofluorescence and HCR fluorescent in situ hybridization showing co-expression of aPKC and *Tbx5* gene expression. Note the low levels of aPKC staining and lack of polarized expression in the forelimb field and (**P-R**) LAMININ expression.

### Cell polarity markers are not translationally regulated during EMT

The preceding experiments suggested that EMT occurred between 12 and 13 somites while forelimb-specific transcription (mediated by Tbx5) likely occurred after 14 somites. To determine if EMT components are translationally regulated, we used ribosome profiling by isotachophoresis (Ribo-ITP) (Ozadam et al., 2023), an ultra-low input ribosome profiling technique to isolate ribosome footprints (RPF) from dissected single somatic LPMs. These regions, containing ∼4,000 cells, also included the overlying ectoderm (Fig. 3A). The resulting data had key characteristics typical of high-quality ribosome occupancy measurements. In particular, RPFs were predominantly 28 nucleotides, which is the length of mRNA protected by a translating ribosome (Fig. 3B). Over 90% of the RPFs mapped to the coding regions of transcripts and showed enrichment around the translation start and stop sites (Fig. 3C, D).

**Figure 3:**
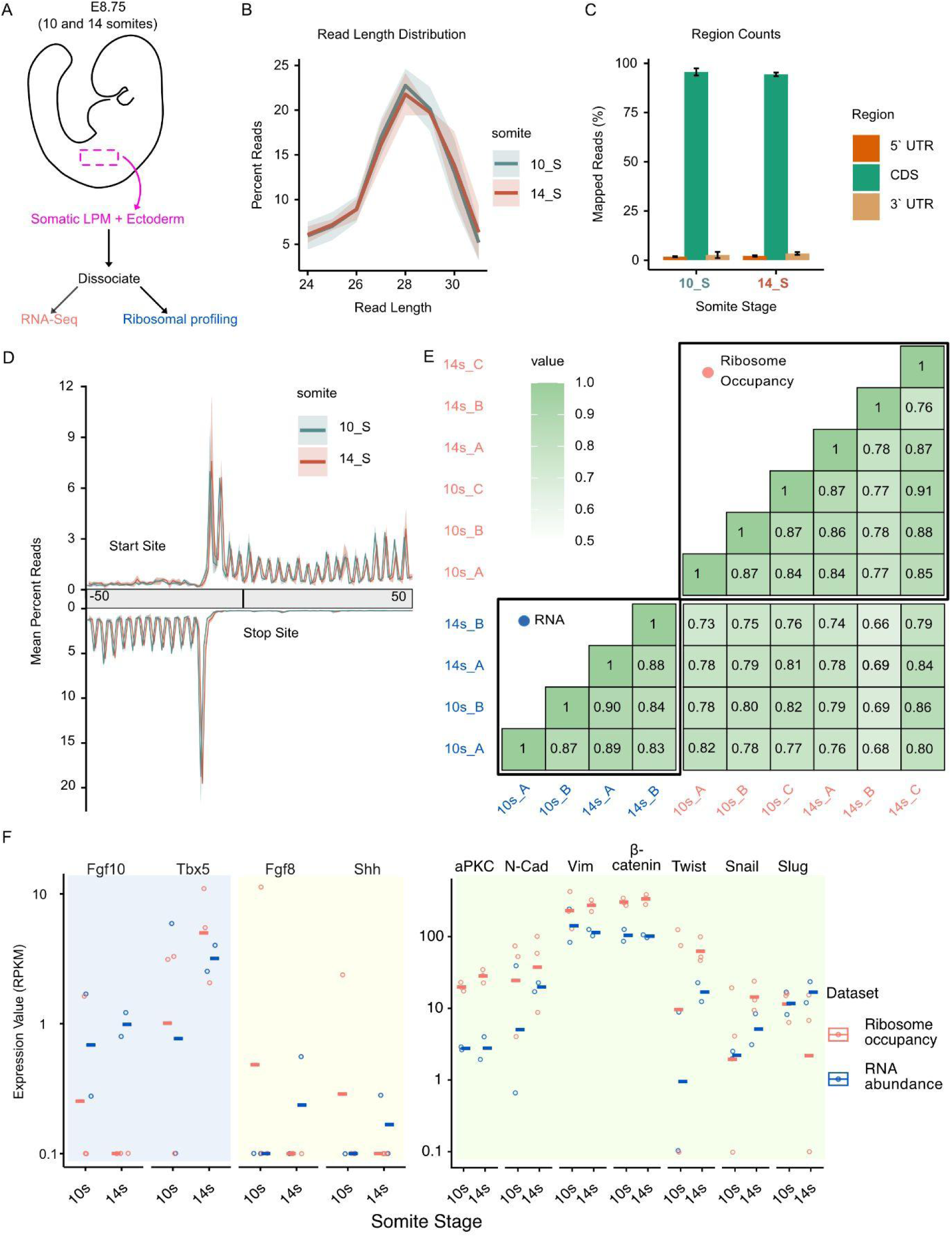
EMT markers are robustly translated in early the limb bud during 10 and 14 somite stages. (A) Schematic of the experiment showing the dissected region of the embryo used to perform paired RNA-Seq and Ribosome profiling. (B) Length distribution of ribosome protected footprints for the 10 and 14 somite stages, shaded area shows deviation between biological triplicates. (C) Mapping of ribosome profiling reads to different transcript features (CDS, 5′ UTR, and 3′ UTR) is presented for each stage. Error bars indicate the standard deviation. (D) Metagene plots for ribosome occupancy around the translational start (top) and stop (bottom). Note that the 5’ ends of mapped reads are plotted hence the characteristic offset relative to the start and stop site are observed. (E) Pairwise correlation matrix of the generated RNA-Seq and ribosome occupancy libraries. Boxes are color coded with darker shades representing higher Spearman’s correlation coefficient. (F) RNA expression and ribosome occupancy of limb bud specifier genes (*Fgf10, Tbx5*), genes not detected in limb bud (*Fgf8, Shh*) and cell polarity and EMT related genes: aPKC (*Prkci),* N-Cadherin *(Cdh2),* Vimentin *(Vim),* β-catenin *(Ctnnb1)*, Twist *(Twist1)*, Snail*(Snai1)* and Slug *(Snai2)* at 10 and 14 somite stages. RPKM values (y-axis) for each biological replicate are shown in circles with the mean depicted with a horizontal line.

We next sought to identify the translational landscape of the early developing limb bud. The Spearman correlation coefficients for pairwise comparison between the ribosome profiling libraries ranged from 0.77-0.9 (Fig. 3E). We also measured RNA abundance using paired RNA-seq from a portion of the same samples, where the correlation coefficients ranged from 0.83 to 0.9. The correlation between the RNAseq and ribosome profiling libraries was slightly lower (0.66-0.86). We observed that replicates from the same stage were as similar to one another as they were to samples from different stages for both RNA-seq and ribosome profiling datasets when using Spearman correlation of transcript abundance. This suggested that the biological variability of ribosome occupancy profiles between 10s and 14s stage limb buds is of similar magnitude to the variation between replicates of the same stage. It also suggested that there are likely limited changes between 10-14 somites consistent with our prediction that this precedes forelimb-specific transcription.

The ribosome occupancy measurements also enabled us to examine the translation status of transcripts involved in cell polarity and EMT. Consistent with its spatial expression (Fig. 1A-D), *Tbx5* was expressed at low levels at the 10-somite-stage and at higher levels in 14-somite-stage limb buds (Fig. 3F, left panel, Dataset S1). *Fgf10* was present at both stages at low levels (Fig. 3F, left panel, Dataset S1). In contrast, *Fgf8* and *Shh*, which are not expressed at these stages, did not have detectable RNA abundance and ribosome occupancy (Fig. 3F, middle panel, Dataset S1). Multiple cell polarity markers were expressed and robustly translated at both of these stages including aPKC (*Prkci*), N-Cadherin (*Cdh2*), Vimentin (*Vim*) and β-catenin (*Ctnnb1*) as well as EMT specific transcription factors Twist (*Twist1*), Snail (*Snai1*) and Slug (*Snai2*) (Fig. 3F, right panel, Dataset S1). These observations suggest a lack of large magnitude changes in the translational efficiency of proteins involved in limb bud initiation or EMT. Notably, we did not detect significant changes in the translational efficiency of transcripts encoding N-Cadherin or β-catenin during the time where they lose their apical polarity (Fig. 2A-C, G-I).

### Single cell RNA sequencing (scRNA-seq) reveals early onset marker genes in the forelimb

The preceding experiments suggest that the onset of TBX5-mediated forelimb specification occurs between 14 and 16 somites. To identify early forelimb-specific genes, we performed scRNA-seq using WT and *Fgf10^-/-^* embryos at 15-16 somites (n=2 per genotype). Embryos were transversely bisected at the axial level of the forelimb (between 7 and 13 somites; Fig. 4A) and libraries were generated using a split pool barcoding method. These libraries included the somatic LPM as well as all other embryonic tissues present at this axial level. Initial clustering did not reveal any genotype specific differences in gene expression profiles or cell-type proportions across clusters, consistent with Fgf10 having a slightly later role in limb development (Fig. S3, Dataset S2). There also were no significant changes in genotype specific gene expression (avg Log2FC > 2, p_adj_ < 0.05) in the somatic LPM cluster (Dataset S3). In all subsequent analyses, the four samples were pooled together. Using marker gene expression, we identified 13 distinct cell populations that represented most embryonic tissues (Fig. 4B, Fig. S4A-K). These included a population of LPM that contained enrichment of splanchnic as well as somatic LPM markers (Fig. 4C). After subsetting and reclustering, these cells could be grouped into three populations (Fig. 4D). We labelled the *Tbx5*, *Prrx1* positive cluster as somatic LPM, the *Fendrr, Hand1* positive cluster as splanchnic LPM while a third, smaller population was referred to as other LPM (Fig. 4D,E). These populations contained many of the same genes identified as splanchnic and somatic LPM-specific in chick embryos (Newton et al., 2022) (Dataset S4). For example, the top marker genes (most differentially expressed in a given cluster) for the somatic LPM included *Tbx5*, *Fgf10* and *Prrx1*, consistent with expectations for this population. However many other top marker genes lack known functions in limb development. Overall, these early onset genes suggest the activation of previously unknown processes in the nascent forelimb.

**Figure 4:**
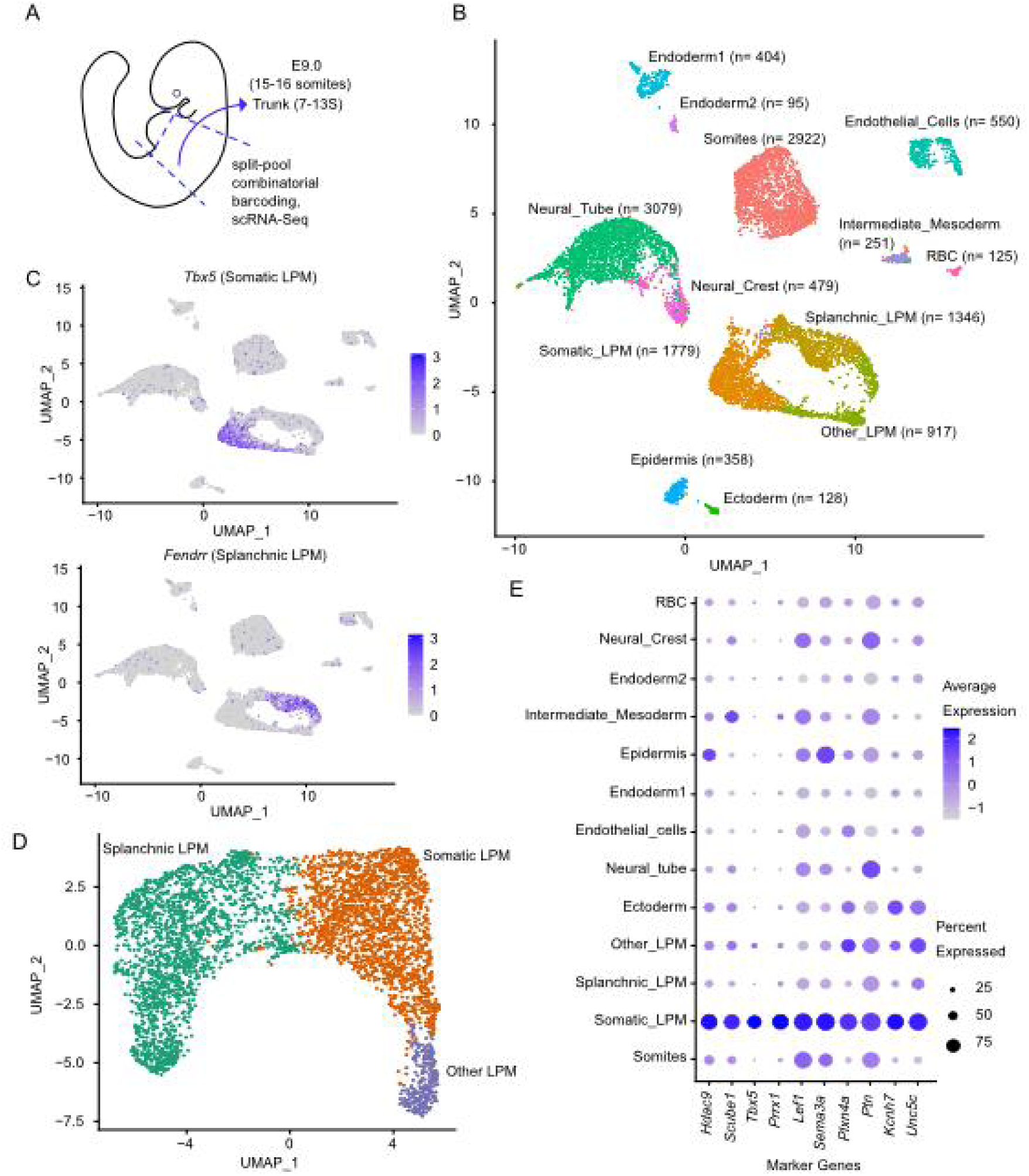
scRNA sequencing of early limb buds. (A) Schematic showing the dissected region of 15-16 somite stage embryos used for split-pool barcoding based single cell RNA sequencing. (B) UMAP plot showing the 13 different cell populations identified in and around the developing mouse limb bud. (C) UMAP plots showing expression of the top somatic LPM and splanchnic LPM cluster marker. The color indicates average expression in a cell. (D) UMAP plot after reclustering the LPM clusters. (E) Dotplot showing the expression of top 10 somatic LPM cluster markers from D across the 12 clusters shown in B. Size of the dot indicates percent cells expressing the gene and color indicates average expression across cells.

We next asked if these early onset genes provide increased in-dataset predictive power in identifying somatic LPM cells. We performed random forest classification of cell clusters based on the individual expression of each of the top 10 marker genes. Among these, *Tbx5* was the best single predictor of somatic LPM cells (Fig. 5A, orange) compared to the remaining nine genes (Fig. 5A, orange). However, combining the expression of *Tbx5* with the other top 10 genes further increased the prediction accuracy (Fig. 5A, blue). In particular, this suggests that expression of *Kcnh7*, *Sema3a*, *Unc5c*, *Plxna4* and *Scube1* could help identify somatic LPM cells during the presumptive limb bud formation stage. We next examined the pairwise expression of *Tbx5* with *Sema3a*, *Unc5c*, *Plxna4* and *Scube1* in somatic LPM and other cells. Cells co-expressing any of these genes with *Tbx5* were more likely to be somatic LPM cells as suggested by the random forest classification (Fig. 5B).

**Figure 5:**
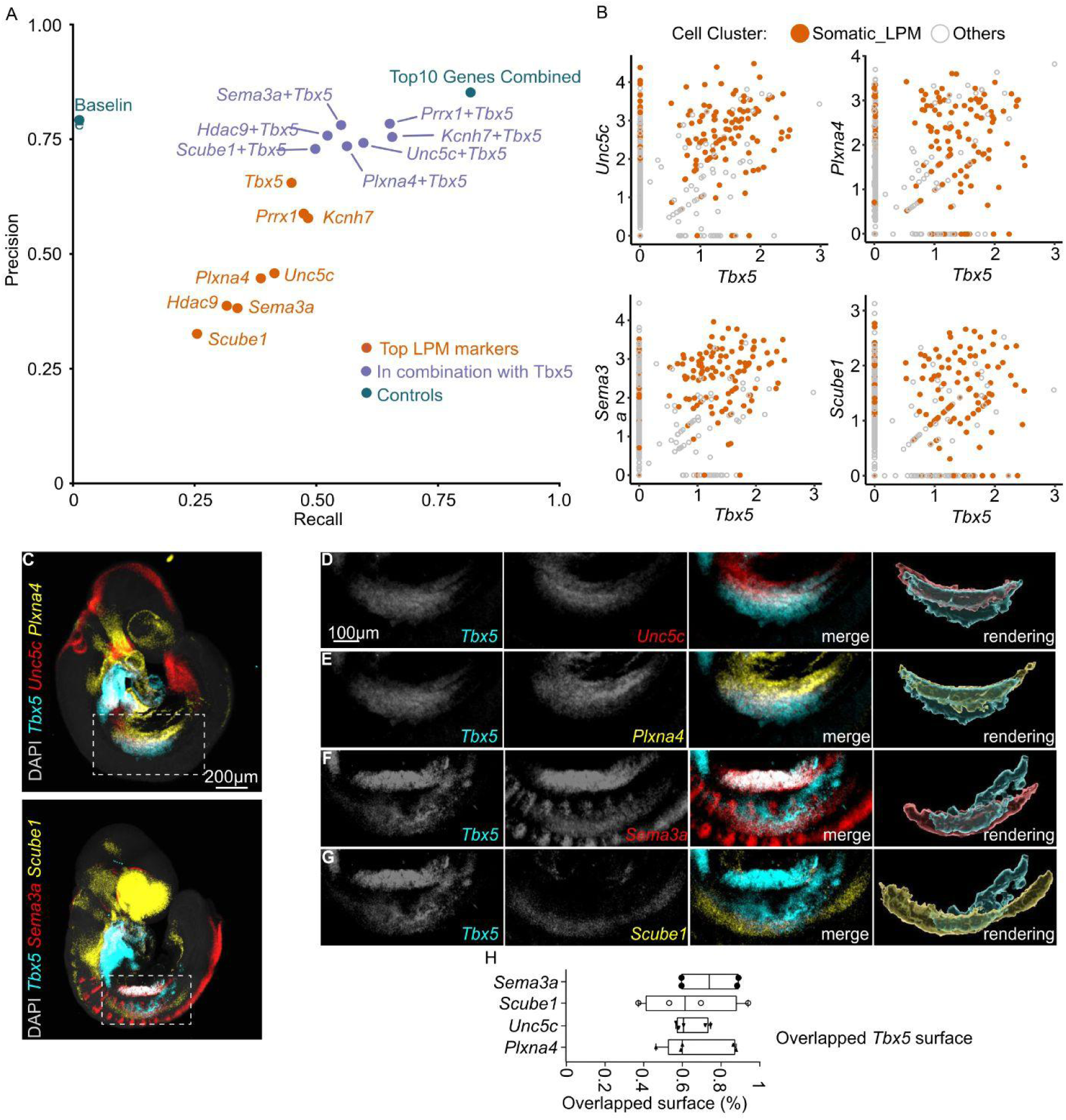
Signature genes improve identification of somatic LPM. (A) Precision recall plot for the random forest predictions using the top 10 marker genes in predicting somatic LPM cells. Marker genes alone (orange) or in combination with Tbx5 (blue) and probability of random prediction (Baseline) and all top10 genes together (cyan) are shown. Precision is the proportion of number of times the prediction was correct and recall is the proportion of somatic LPM cells that were correctly predicted by the gene. (B) Overlaid HCR image showing the expression of *Tbx5*, *Unc5c*, *Plxn4a* and *Tbx5*, *Sema3a*, *Scube1* in 15-16 somite whole mouse embryos, (C) Correlation plots showing the expression of Tbx5 with the experimentally validated marker genes (*Unc5c*, *Plxna4*, *Sema3a* and *Scube1*). 1000 random cells are plotted for effective visualization. Log-normalized expression values are plotted on both the x-axis and y-axis. (D-G) Zoomed images of the developing limb field in 15-16 somite embryos. The images on the far right depict volumetric renderings showing the overlap between the *Tbx5* expression domain and that of the indicated gene in the forelimb field (H) Quantification of the overlap between the *Tbx5* domain and the indicated genes. (D-H) n=4 biological replicates for *Sema3a* and *Scube1* and n=5 for *Unc5c* and *Plxna4*.

Given that the predictive performance was evaluated without embryo-stratified cross-validation; these results reflect in-dataset behavior and may overestimate generalization to new embryos. Hence, we examined the spatial expression of five of these genes in an independent set of 15-16-somite-stage wild-type embryos using HCR fluorescent in situ hybridization. We failed to detect any embryonic expression of *Kcnh7*, likely due to probe failure. However, the other four genes showed expression in the nascent forelimbs (Fig. 5C-H), including substantial overlap with *Tbx5* (Fig. 5D-H), confirming the identification of these genes as markers of the nascent forelimb.

### Identification of early onset TBX5 target genes in the forelimb

The identification and subsequent validation of early marker genes in the 15-16 somite somatic LPM suggested that they are likely to be among the first genes expressed in the nascent limb bud and as such may mediate key processes underlying TBX5-mediated forelimb specification. The top 30 marker genes set was significantly enriched for genes involved in neuron projection and cell adhesion, suggesting their potential involvement in early forelimb formation (Figure 6A). We hypothesized that they might also represent early direct targets of TBX5-mediated transcription. To test this, we identified TBX5 binding regions by ChIP-seq, using E10.5 forelimbs because of the challenges in obtaining enough tissue at 15-16 somites. We identified a total of 13,581 binding regions (FDR<0.05). Of these, 6957 were associated with 4776 genes using GREAT (Abler et al., 2009; McLean et al., 2010) (Dataset S5). Consistent with our hypothesis, the top 30 marker genes from the scRNA-seq analysis were substantially enriched for TBX5 binding regions and also contained significantly more TBX5 binding regions per gene compared with the remaining TBX5-associated genes (Fig. 6B-D; Dataset S6). These regions included *Tbx5* itself, consistent with its proposed role in autoregulation (Sun et al., 2004) and *Fgf10*, a known target gene (Agarwal et al., 2003; Delgado et al., 2021; Rallis et al., 2003). They also included *Sema3a*, *Plxna4*, and *Unc5c* (Fig. 6E-H) which are all expressed in the forelimb at 15-16 somites (Fig. 5D-H) as well as *Scube1*. While these regions likely represent immediate early targets of TBX5-dependent transcriptional programs in forelimb initiation, this interpretation carries certain caveats as discussed below.

**Figure 6:**
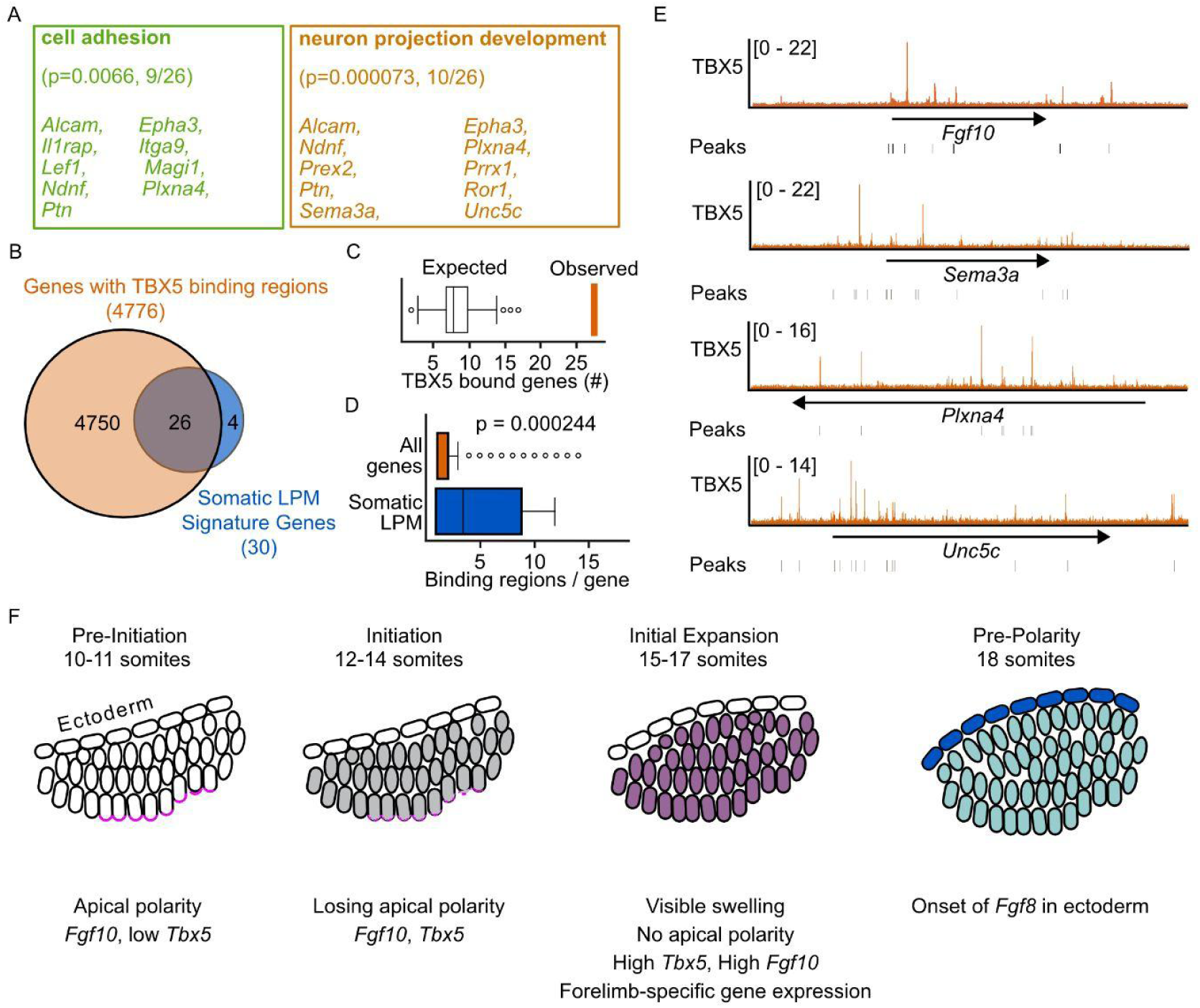
Somatic LPM signature genes are TBX5 targets. (A) The TBX5 signature target gene set is significantly enriched for cell adhesion and neuron projection development GO terms (terms indicate the adjusted p-value and number of enriched genes. (B) Vein diagram showing intersection between all genes with TBX5 binding regions and somatic LPM signature genes identified by ChIP sequencing for TBX5 from E10.5 stage embryos. (C) The expected versus observed number of genes with TBX5 binding sites. The number of TBX5-associated genes observed in the Somatic Lateral Plate Mesoderm group (26/30) was compared to a null distribution generated from 10,000 random draws of 30 genes. (D) Box plot showing the number of TBX5 binding regions per gene for somatic LPM signature genes and all other genes. (E-H) Coverage plots showing TBX5 ChIP-seq reads across *Fgf10*, *Sema3a*, *Plxna4* and *Unc5c* genomic regions respectively. (I) Graphical illustration of the phenotypic and gene expression changes happening in the somatic LPM from 10-18 somites.

## DISCUSSION

Here, we describe the early stages of mouse forelimb specification in the context of marker gene expression, cell polarity and sequencing based high-throughput approaches (RNA-seq, scRNA-seq, ChIP-seq, and Ribo-ITP). These include the temporal expression of genes initiating forelimb development, the occurrence of EMT, and the onset of forelimb morphology and gene expression. These events have been individually described, but many were known from earlier studies, often performed decades ago, that did not focus on the temporal onset or how they compared with other hallmarks of forelimb development. Building on this framework, we focused on genes expressed at the onset of forelimb initiation. The loci around these genes are highly enriched for TBX5 binding sites, suggesting that they may represent transcriptional targets of TBX5. Many of these genes encode for proteins involved in neuron projection and cell adhesion, identifying candidate genes and processes for TBX5-mediated forelimb specification.

We propose that forelimb initiation is organized into four distinct phases (Fig. 6F). The Pre-Initiation stage (10-11 somites) consists of multilayered somatic LPM that expresses *Fgf10* and low levels of *Tbx5*. The Initiation stage (12-14 somites) is characterized by the gradual loss of apical polarity markers and a slight increase in the levels of *Tbx5*. N-Cadherin and Beta-Catenin, initially apically enriched in the most lateral layer of somatic LPM, lose this enrichment between 12-13 somites, suggesting that this marks a transition point in limb development that is consistent with the upregulation of *Tbx5*. This is followed by the Initial Expansion stage (15-17 somites), which marks the first visible indication of bud and is characterized by high levels of *Tbx5*, *Fgf10* and the expression of downstream target genes. The Pre-Polarity stage (starting at 18 somites) is defined by the gradual onset of ectodermal *Fgf8*, which will later activate proximal distal polarity (Sedas Perez et al., 2023). It is important to emphasize that the limb bud will continue to express uniform expression of *Meis1*, a readout of proximal-distal polarity within the mesenchyme up until 24 somites, suggesting that the limb bud mesenchyme itself remains unpolarized well after this timepoint (Delgado et al., 2020). We also note that *Fgf8* is itself downstream of several upstream ectodermal regulators, including Sp6/Sp8 and Wnt3/Beta catenin (Barrow et al., 2003; Haro et al., 2014; Soshnikova et al., 2003), indicating the initial ectodermal response to mesenchymal signaling precedes this timepoint. For these reasons, we utilized 15-16 somite samples to identify early forelimb-specific target genes that were less likely to be influenced by ectodermal feedback.

It is notable that *Ffg10* is robustly expressed throughout the somatic LPM many stages prior to the upregulation of *Tbx5*. Early *Fgf10* does not regulate *Tbx5* expression domains during the Initiation or Initial Expansion phases (Fig. S1) nor does it detectably alter gene expression (Dataset S2). These observations are consistent with the known role of Fgf10 being genetically upstream of *Fgf8* (Ohuchi et al., 1997) and suggest that Fgf10 does not have additional roles in regulating gene expression within the early forelimb mesenchyme. We interpret this to mean that TBX5 likely maintains and upregulates Fgf10 in the nascent forelimb after specification rather than initiating its expression.

The elimination of apically polarized N-Cadherin in the most medial somatic LPM provides a clear endpoint for EMT. However, in the absence of live cell imaging, we are not able to determine when this process first initiates. The ribosomal profiling experiments suggest that this is not triggered by changes in the translational efficiency of *Snail* or *Twist.* The somatic LPM consists of a single layer at 7-8 somites (Fig. S2A-D) and is multi-layered by 10 somites. Since the more lateral layers of cells do not exhibit polarity, the process of losing mesenchymal polarity could be a gradual process, perhaps occurring when daughter cells move more laterally. Both *Tbx5* and *Fgf10* promote EMT (Gros and Tabin, 2014). Given that Tbx5 is not upregulated until 14 somites, after EMT is largely completed, it remains unclear how this might occur, although we cannot rule out the possibility the lower levels of TBX5 present at 12-13 somites might be sufficient for this process.

The somatic LPM cells that are present between the ectoderm and the medial layer of somatic LPM lining the coelom are likely to have different adhesion properties based on their differential expression of polarized cell markers. For example, the lateral somatic LPM cells might be more motile than their polarized medial counterparts. Since somatic LPM cells have been shown to exhibit TBX5-mediated directional migration in forming the zebrafish pectoral fin and exhibit coordinated movement into the nascent mouse limb bud (Boyle-Anderson et al., 2022; Mao et al., 2015; Wyngaarden et al., 2010), the breakdown of polarized mesenchyme could enable the additional motility required to initiate the forelimb bud. FGF10 could also play a role in regulating this motility in a similar fashion to how FGF signaling regulates cell motility in the presomitic mesoderm and in the distal limb bud (Bénazéraf et al., 2010; Gros et al., 2010).

We used scRNA-seq on embryos dissected at the axial level of the forelimb field to define 30 signature genes in the Expansion phase. The dataset, which we subsequently validated by gene expression, likely includes the first genes expressed in the nascent forelimb. These genes are highly enriched for TBX5 binding regions, which are present in 26/30 genes (Fig. 6A-C), suggesting that they represent direct TBX5 target genes. There are two caveats to this assumption. First, the TBX5 ChIP-seq data was collected on E10.5 forelimbs rather than 15-16 somite (∼E8.75) somatic LPM. While beyond the scope of the current manuscript, future work implementing low input binding assays for TBX5 at these early stages would provide direct evidence of binding at this timepoint. Second, we did not determine if these genes were downregulated in *Tbx5^-/-^* limb buds, as we reasoned that genes within this domain were unlikely to be expressed in the complete absence of a limb bud that occurs in *Tbx5^-/-^* embryos. Nonetheless, these early onset genes are particularly interesting since TBX5 has a transient role in initiating forelimb outgrowth (Hasson et al., 2007). In addition to including the known TBX5 target gene, *Fgf10* (Agarwal et al., 2003; Delgado et al., 2021; Rallis et al., 2003), the gene set includes several other genes involved in somatic LPM development or limb specification. Notably, it is enriched for genes involved in neuron projection as well as cell adhesion (Fig. 6A).

The presence of neuron projection factors, including *Plxna4*, *Sema3a*, *Unc5c* and *Epha3* indicates that the nascent forelimb initially expresses multiple genes thought to be involved in guiding the axons that will later project to the limb (Kania and Jessell, 2003; Noguchi et al., 2017; Osterwalder et al., 2014; Poliak et al., 2015; Suto et al., 2005; Vaidya et al., 2003). However, axonal projection to the forelimb bud does not initiate until after E11, days after limb bud initiation and the acquisition of later limb polarity signals. On the other hand many of these factors are also implicated in cell adhesion. The enrichment of cell adhesion factors suggests that they might underlie TBX5-mediated initiation of forelimb outgrowth, a scenario that was first proposed by Ho and colleagues (Ahn et al., 2002). Classical studies have shown that limb bud cells have differential cohesiveness compared to flank tissues and suggested that cohesiveness plays a role in limb bud generation (Damon et al., 2008; Heintzelman et al., 1978; Wada, 2011). This role could include impacting TBX5-directed migration if this turns out to be similar in mouse embryos to that observed zebrafish pectoral fin formation (Boyle-Anderson et al., 2022; Mao et al., 2015). Morphogenesis in the established early limb bud (21 somites) occurs by durotaxis, where differential mesodermal stiffness regulates the shape of the limb bud (Zhu et al., 2024) and it is possible that changes in adhesion might catalyze this process.

## METHODS

### Mouse embryos

Experiments involving mice were approved by the Institutional Animal Care and Use Committee at the University of Texas at Austin (AUP-2022-00221) or the University of Georgia. Embryos were obtained from Swiss Webster mice with the exception of Fgf10^+/-^ experiments, which used the Fgf10^tm1.1Sms^ allele (Abler et al., 2009) maintained on a mixed background (∼75% Swiss Webster).

### Wholemount In Situ Hybridization

For colorimetric wholemount in situ hybridization, embryos were treated with 10 μg/mL proteinase K (Invitrogen 25530015) for 3 min, pre-hybridized at 68C for 1 h and hybridized with 500 ng/mL digoxigenin-labeled antisense RNA at 68C overnight and developed with BM purple (Sigma-Aldrich 11442074001). Embryos were postfixed and cleared in 100% glycerol prior to imaging on a Leica M165 stereomicroscope with a DFC420C Leica camera. Fluorescent HCR 3 in situ hybridization was performed as described previously (Anderson et al., 2020) using 16nM of each probeset (Molecular Instruments). The probesets included Tbx5-B2 (lot RTB930), Sema3a-B1 (lot RTN635), Scube1-B3 (lot RTT135), Unc5c (lot RTB893) and Plxna4-B3 (lot RTT770). Cleared samples were imaged every 20µm on a Nikon AXR confocal microscope with a plan apo 4X objective and post-processed in Imaris.

### Immunofluorescence and multiplexed HCR in situ hybridization

Cryosections were hybridized overnight at 37C with a 4nM *Tbx5*-B4 probe solution and then amplified for 4-24 hours using RNA FISH buffers with B4-Alexa Fluor 647 hairpins (Molecular Instruments) according to the vendor’s protocol: HCR™RNA-FISH(v3.0) protocol for fresh frozen or fixed frozen tissue sections). After amplification, the slides were washed 3 times over an hour in 5XSSC with 0.1%Tween 20, 3×5min with PBS, blocked for 1 hour at room temperature (5% Normal Goat Serum, 3% BSA, 0.1%Tween 20 and incubated with primary antibodies overnight at 4C. The next day, samples were intubated in secondary for 1 hour, DAPI 1:5000 for 5 minutes and mounted with Prolong Gold. They were imaged on a Zeiss LSM 710 confocal microscope The following antibodies were used: N-Cadherin (1:200, Cell Signaling Technology CST13165 Lot# 4); aPKC (1:100, Santa Cruz BIotechnology SC-17781 Lot#C1122); Laminin (1:100, Sigma-Aldrich L9393 Lot#0000201695); Beta-Catenin (1:100, Santa Cruz Biotechnology SC-7963 Lot#K1022); Twist (1:100, Abcam ab50887 Lot# GR3351419-9). Alexa Fluor 488-conjugated Goat anti-Rabbit IgG (1:500, ThermoFisher A11034 Lot# 238003); Alexa Fluor 568-conjugated Goat anti-Mouse IgG (1:500 ThermoFisher A11004 Lot# 2447869).

### RNA-seq and Ribo-ITP of microdissected mouse limb bud

10-and 14-somite embryos were bisected transversely between 7-and 10-somites in dPBS. These core regions were transferred to a base Ribo-ITP buffer containing 20 mM Bis-Tris, 5 mM MgCl2, 5 mM CaCl2, 100 mM NaCl, 1 mM DTT, 0.25% BSA and the somatic LPM and overlying ectoderm was removed from the left side of the embryo using dissecting knives and flash frozen. Frozen limb bud tissue samples were thawed on ice and Triton-X 100 and Cycloheximide were added to a final concentration of 0.1% and 0.1 mg/mL respectively. The lysate was separated into two tubes, 5μL for Ribo-ITP and 3μL for SmartSeq3 (Hagemann-Jensen et al., 2020). For Ribo-ITP, 1μL 1:300 diluted RNaseI (Ambion) was added and digestion was performed at 37°C for 30mins. Digestion was halted by adding 1μL of 0.7% SDS following which Ribo-ITP was performed as described in (Ozadam et al., 2023). The final indexing PCR was performed for 15 cycles. SmartSeq3 was performed as described in (Ref) with the following modifications, the preamplification PCR was performed for 13 cycles and indexing PCR was performed for 16 cycles. Both the ribosome profiling and RNASeq libraries were cleaned with AmPureXP beads with additional PAGE purification for Ribo-ITP libraries. The libraries were sequenced by Illumina Novaseq 6000.

### Analysis of ribosome profiling and RNA-Seq data

The ribosome profiling libraries were processed using the Riboflow pipeline v.0.0.1 using deduplication with unique molecular identifiers - UMItools (Ozadam et al., 2020). The RiboR package was used for making metagene plots, region coverage plots and read length distribution and RiboGraph for data QC and visualization (Chacko et al., 2024). Ribosome profiling reads with read length between 26-31nt were selected. These raw read counts were then RPKM-normalized using edgeR v4.2.2 (Robinson et al., 2009) with the transcript length. Normalized reads were used for calculating Spearman Correlation coefficients and plotting gene expression values.

### scRNA-seq

Two 15 somite-stage Fgf10^+/+^ and two 16 somite-stage Fgf10^-/-^ embryos (E8.75) were dissected in cold 1X DPBS. The heart and tissues anterior to somite 7 and posterior to somite 13 were dissected away. The resulting core of embryonic tissue included the forelimb field as well as other embryonic tissues present at this axial level. Single embryos were trypsinized and then filtered through a cell strainer, transferred into a 1% BSA coated tube and centrifuged for 10 minutes at 500g at 4°C. Cells were fixed as described using (Tran et al., 2022) and ∼58-73k cells per embryo were stored in-80℃. scRNASeq libraries were prepared using the Parse Evercode WT Mini v2 kit according to the manufacturers protocol. The libraries were sequenced on an Illumina NovaSeq X Plus. A total of 14,214 cells were sequenced across all the limb buds with a depth of 32,104 reads/ cell. We detected 7034 median transcripts/cell and 3,188 genes/ cell.

### Analysis of scRNAseq data

The scRNA-seq datasets were analyzed and visualized using the Parse Biosciences Pipeline Trailmaker^TM^ web tool (Pipeline v1.2.1, analysis completed on 09.16.2024). The raw FASTQ files were processed using Trailmaker^TM^ pipeline module to demultiplex the combinatorial barcodes and obtain cell-by-gene count matrices. These matrices were then integrated into the Trailmaker^TM^ Insights module for further analysis where the barcodes were filtered through four sequential steps for filtering cells based on # of barcodes per cell, removing dead cells by filtering for cells with high mitochondrial gene content, filtering for number of genes vs transcripts and removing doublet cells using automatic filtering for each sample. Post filtration, data was log-normalized, and the top 1000 highly variable genes were selected using the variance stabilizing transformation method. Principal component analysis was performed, and the top 25 principal components, explaining ∼80% of the total variance were used for batch correction with Harmony (Korsunsky et al., 2019). Clustering was performed using Seurat’s implementation of the Louvain method (Blondel et al., 2008). For visualization, a uniform manifold approximation and projection (UMAP) embedding was calculated using Seurat’s wrapper for the UMAP package with cosine distance 0.3. Cluster-specific marker genes were identified by comparing cells of each cluster to all other cells. Ribosomal, mitochondrial and cell cycle genes were excluded from this analysis. Top cluster markers were manually assessed to assign cluster labels. The resulting R object was used for further analysis locally using Seurat v5.3.0. To reduce bias introduced by the sex of the embryos, a list of X and Y chromosome genes was downloaded from GENCODE (GRCm39, v112) and used to filter the features in the Seurat object. This filtered Seurat object was used for all further analyses. Cluster markers for all the cell clusters were identified by comparing the expression across a given cluster and all other clusters.

The three LPM clusters were subsetted. Principal component analysis was performed, and the top 20 principal components, explaining 86.67% of the total variance were retained. Clustering was performed using Seurat’s implementation of the Louvain method, followed by UMAP embedding using Seurat’s wrapper for the UMAP package with cosine distance 0.1. Cluster-specific marker genes were identified by comparing cells of each cluster to all other cells using the RPresto v1.4.7 package. Top 30 marker genes for all the clusters were used for further analysis.

### Random forest prediction of LPM identity based on RNA expression

The top 10 Somatic LPM marker genes with most significant adjusted p-values were selected from the subsetted clustering analysis. To test if these genes were good predictors of SPM cells, the identities of all non-SPM clusters were replaced with ‘other LPM’. randomForest v4.7.1.2 package was used to classify cells as ‘SPM’ or ‘other LPM’ based on expression of top 10 marker gene expression alone and in combination with *Tbx5*. All the top 10 genes combined were used as a positive control and the null model predicted ‘SPM’ cells without any genes. The confusion matrices for each prediction were used to calculate Precision and Recall where Precision = True Positive / (True Positive + False Positive) and Recall = True Positive / (True Positive + False Negative). The resulting values were plotted as a Precision/Recall plot.

### ChIP-seq

ChIP-seq was performed as previously described with minor modifications (Infante et al., 2013). Briefly, mouse forelimb buds were isolated from E10.5 mice (outbred ICR; Envigo). Forelimb samples were crosslinked in 1% formaldehyde for 30 minutes and stored at-80°C until further processing of chromatin. Approximately 50 μg of chromatin was sheared using a Diagenode Bioruptor-300 before incubating overnight with a TBX5 polyclonal antibody (R&D Systems #AF5918). The chromatin-antibody mixture was then incubated for 3 hrs on a Protein G Agarose Column (Active Motif, #53039). After eluting antibody-bound DNA, Illumina libraries were prepared using the NEBNext Ultra II Library Prep Kit and amplified for 15 cycles. Sera-Mag SpeedBeads (Cytiva) were used to perform size selection and to remove primer and adapter fragments. Two independent ChIP and Input biological replicates were generated, and libraries were sequenced by the Georgia Genomics and Bioinformatics Core on the NextSeq-550 platform to generate single-end 75 bp reads. Reads were aligned to the mm10 mouse reference, and ChIP-seq peaks were called using the ENCODE Transcription Factor processing pipeline (https://github.com/ENCODE-DCC/chip-seq-pipeline2) (ENCODE Project Consortium., 2012). Highly reproducible (conservative) peak sets were used with an irreproducible discovery rate (IDR) cutoff of 0.05. TBX5 binding regions were associated with genes using GREAT 4.0.4 (McLean et al., 2010; Tanigawa et al., 2022) with the following parameters:: Basal+extension: 5000 bp upstream, 1000 bp downstream, 100000 bp max extension, curated regulatory domains excluded.

### Statistical Analyses and Visualization

Data was filtered and arranged using functions from the: dplyr v1.1.4, tidyr v1.1.3 R packages. The following R packages were used to generate and arrange plots: pheatmap v1.0.13, ggplot2 v3.5.2, cowplot v1.2.0. For single cell data visualization Seurat plotting functions were used. IGV v. was used for the ChIP-seq peak visualization.

## ACKNOWLEDGEMENTS

We thank Dr. Mark Lewandoski for providing the *Fgf10* mouse line and Dr. Benoit Bruneau for providing the *Tbx5* in situ probe. Microscopy was performed at the Center for Biomedical Research Support Microscopy Facility at UT Austin (RRID:SCR_021756). This work was supported in part by the National Institutes of Health R21HD110096 (SV and CC), R35GM150667 (CC), R01HD073151 (SV), R01HD081034 (DM) and the Welch Foundation F-2027-20230405 (CC).

## DATA AVAILABILITY

All sequencing data and the processed files generated in this study have been submitted to the Gene Expression Omnibus under the accession numbers-GSE310800, GSE310825 and GSE309944. All custom scripts used to perform bioinformatics analyses are available at GitHub (https://github.com/Vigh99/limb-bud-initiaiton/tree/main).

---

**Supplemental Figure 1:**
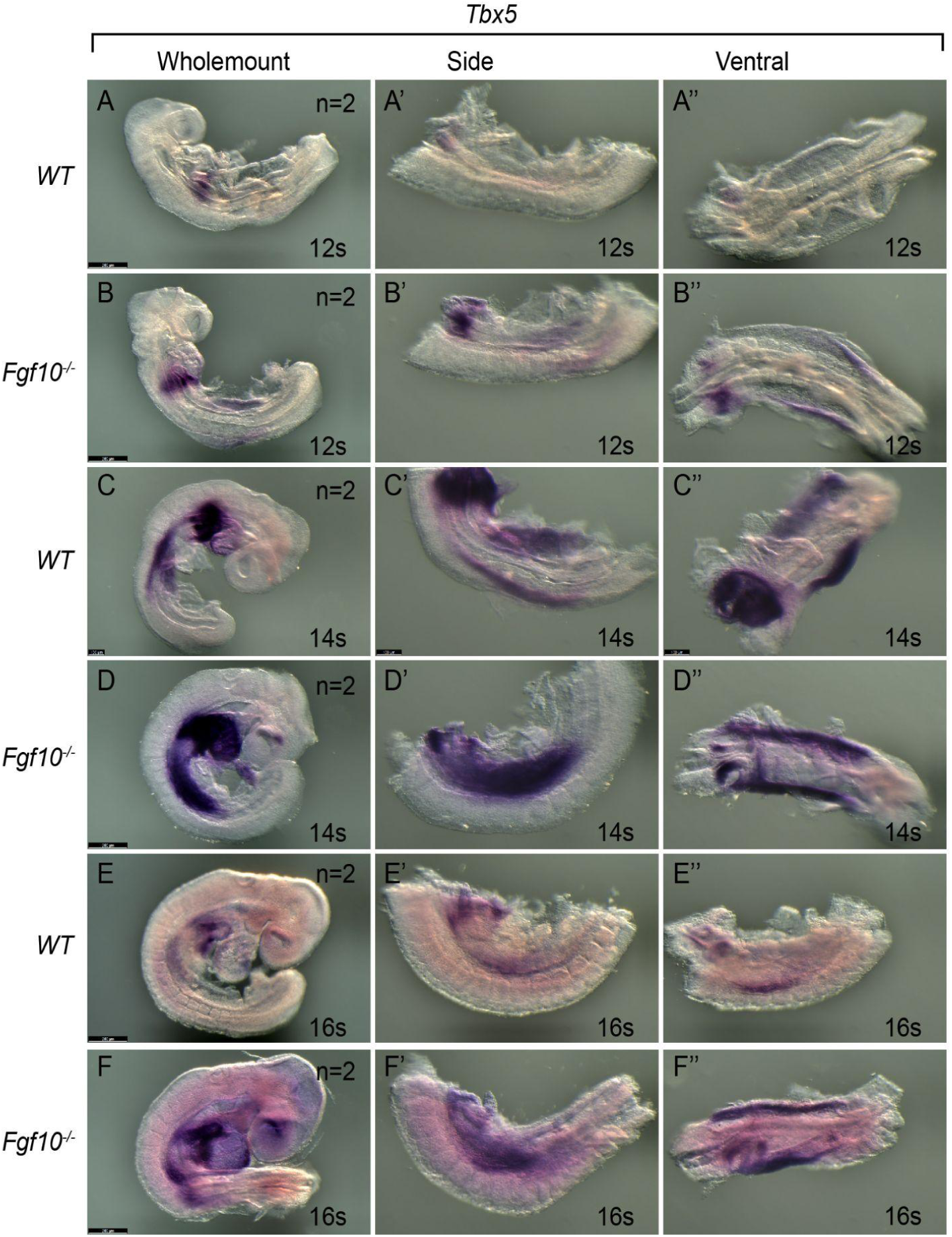
*Tbx5* expression in the forelimb field is unaltered in *Fgf10* loss-of-function mutants. (A-F) Wholemount in situ hybridization on embryos with the somite stage indicated in the bottom right, the genotype indicated on the left, and scale bars at the bottom left. Images in A’-F’’ indicate zoomed-in images of the somatic LPM from the same embryo shown from in A-F from a side (A’-F’) and ventral (A’’-F’’) view. Scale bars indicate 250µm except for panel C, which is 100µm. Each timepoint has 2 replicates per condition.

**Supplemental Figure 2:**
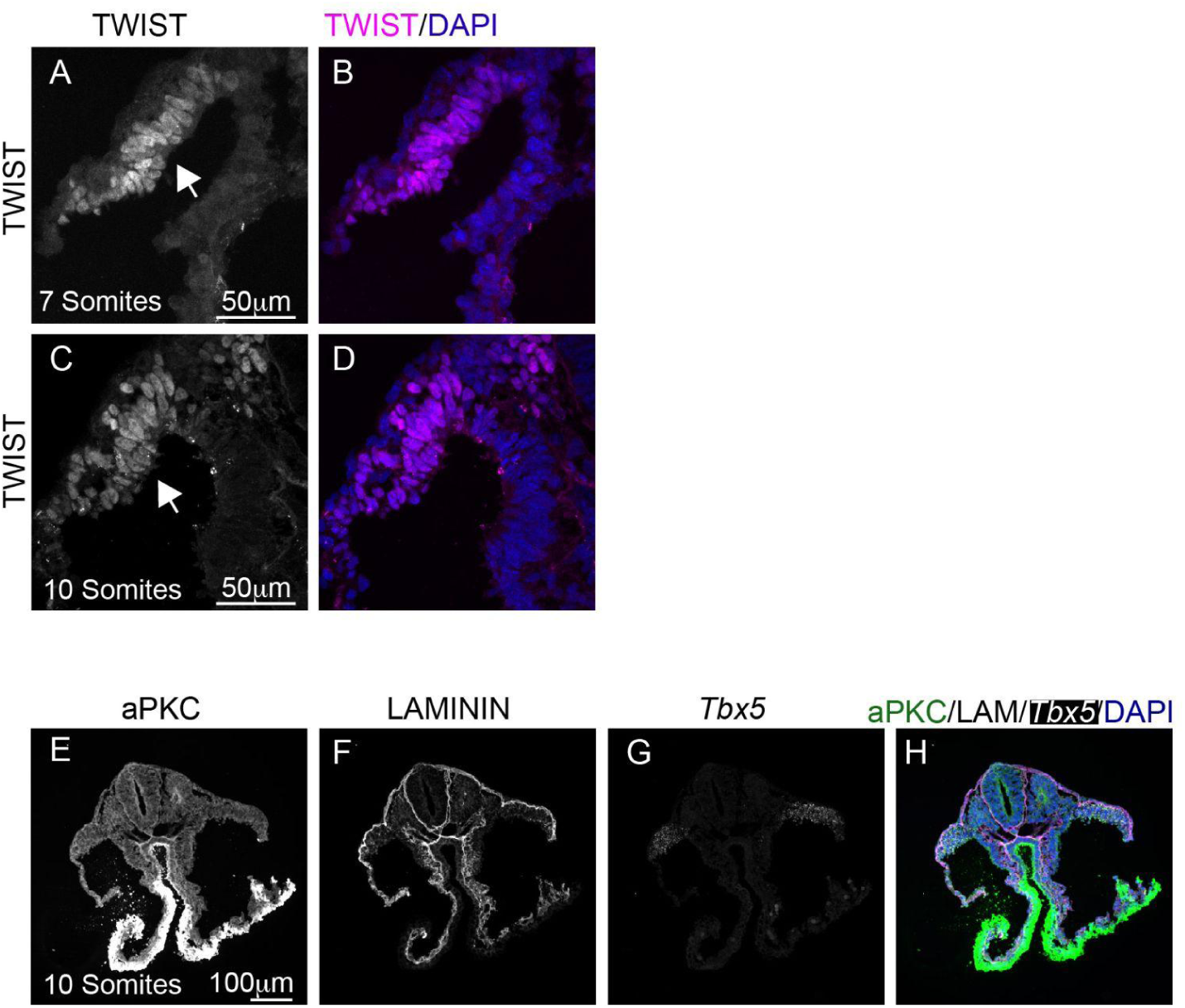
Expression of TWIST and aPKC in the forelimb field. (A-D) Maximal intensity 60X projections showing TWIST expression in the somatic LMP (arrows) at 7-8 somites (n=3) or 10 somite (n=3) embryos in the forelimb field. This antibody recognizes TWIST1 but is likely to cross-react with TWIST2 as well (E-H) Maximal intensity 20X projections showing multiplexed expression of aPKC, LAMININ, and *Tbx5*. This is the same section shown at 60X in Fig 2J-L. The scale bars are indicated within the image.

**Supplemental Figure 3:**
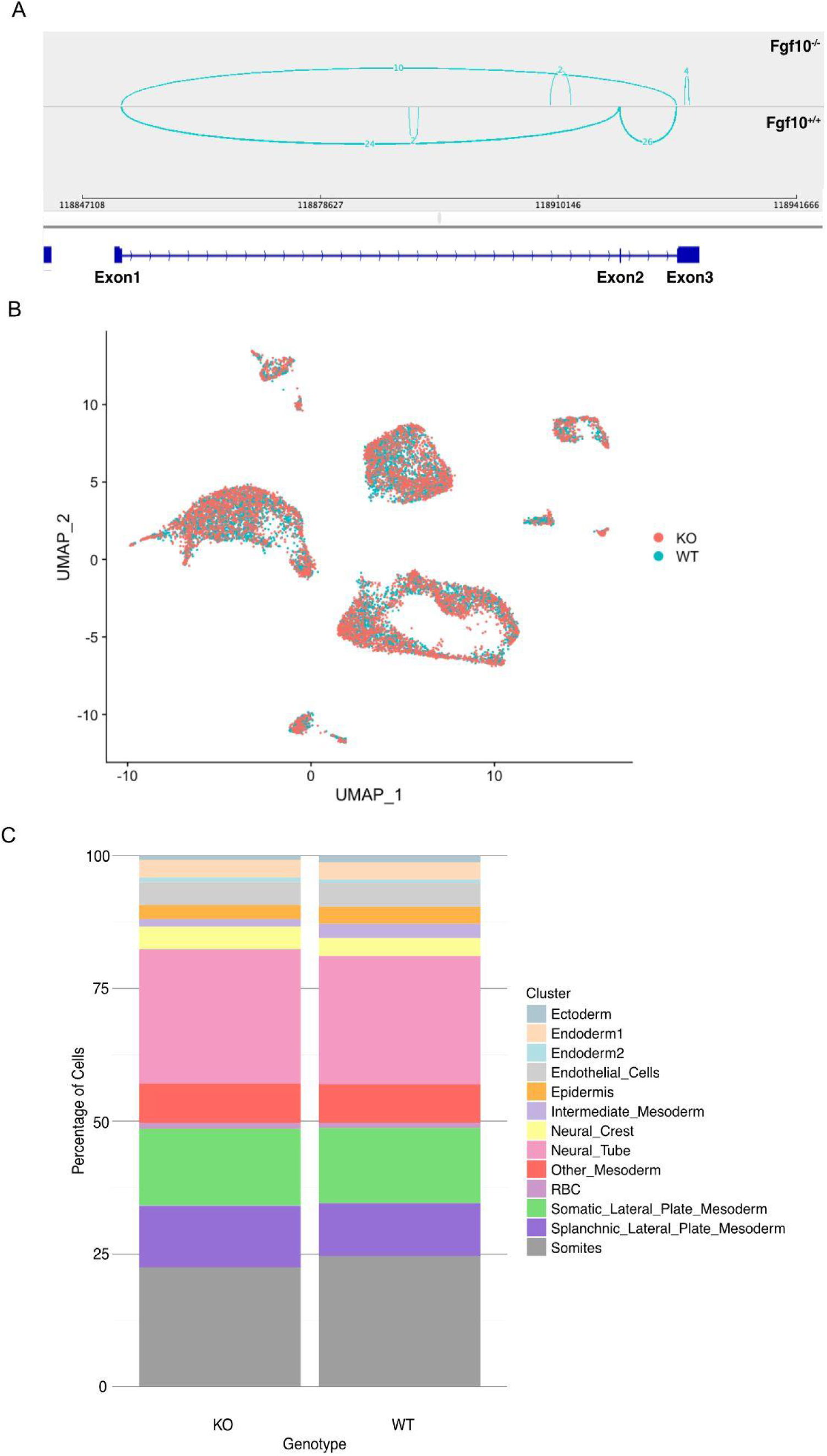
Fgf10^+/+^ v/s Fgf10^-/-^ gene expression. (A) Sashimi plot generated from IGV showing read alignment on the *Fgf10* locus in the *Fgf10**^+/+^***and *Fgf10**^-/-^*** cells. The exons on the *Fgf10* locus are shown as filled blue bars and introns as a blue line, the arrows indicate the direction of transcription. The absence of reads mapping on Exon2 corresponds to the *Fgf10* null allele. (B) UMAP plot showing *Fgf10^+/+^*and *Fgf10^-/-^* cells across the identified clusters. (C) Stacked bar plot showing the percentage of cells in each of the identified clusters in *Fgf10^+/+^* and *Fgf10^-/-^* genotype samples.

**Supplemental Figure 4:**
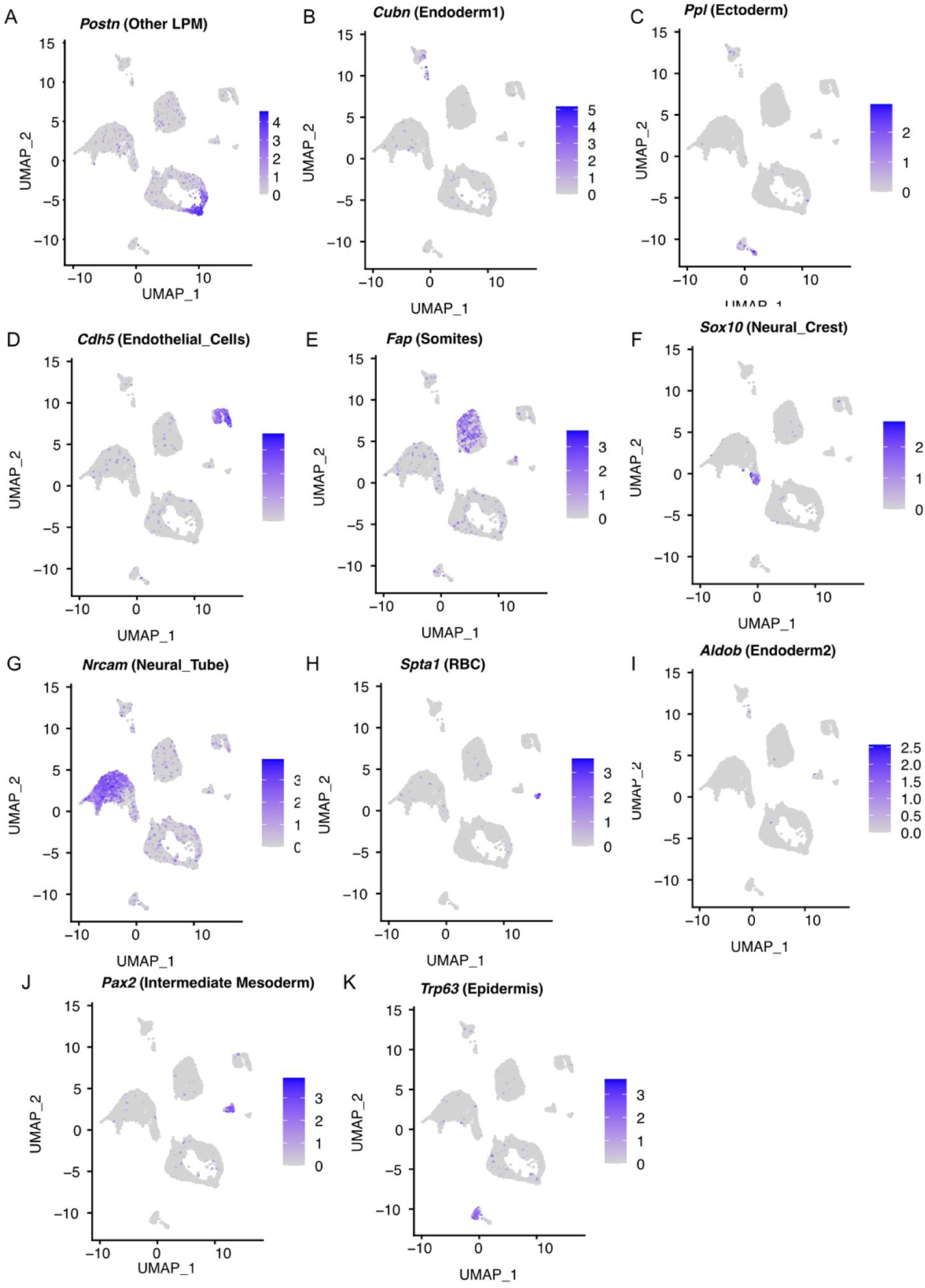
Expression of cluster markers. UMAP plots showing the expression of each cluster marker gene across all the cells sequenced. The name of the cluster are mentioned in parentheses.

## REFERENCES

Abler, L. L., Mansour, S. L. and Sun, X. (2009). Conditional gene inactivation reveals roles for Fgf10 and Fgfr2 in establishing a normal pattern of epithelial branching in the mouse lung. Dev Dyn 238, 1999–2013.

Agarwal, P., Wylie, J. N., Galceran, J., Arkhitko, O., Li, C., Deng, C., Grosschedl, R. and Bruneau, B. G. (2003). Tbx5 is essential for forelimb bud initiation following patterning of the limb field in the mouse embryo. Development 130, 623–633.

Ahn, D.-G., Kourakis, M. J., Rohde, L. A., Silver, L. M. and Ho, R. K. (2002). T-box gene tbx5 is essential for formation of the pectoral limb bud. Nature 417, 754–758.

Anderson, M. J., Magidson, V., Kageyama, R. and Lewandoski, M. (2020). maintains levels critical for normal somite segmentation clock function. Elife 9,.

Barrow, J. R., Thomas, K. R., Boussadia-Zahui, O., Moore, R., Kemler, R., Capecchi, M. R. and McMahon, A. P. (2003). Ectodermal Wnt3/beta-catenin signaling is required for the establishment and maintenance of the apical ectodermal ridge. Genes Dev 17, 394–409.

Bell, S. M., Schreiner, C. M. and Scott, W. J. (1998). The loss of ventral ectoderm identity correlates with the inability to form an AER in the legless hindlimb bud. Mech Dev 74, 41–50.

Bénazéraf, B., Francois, P., Baker, R. E., Denans, N., Little, C. D. and Pourquié, O. (2010). A random cell motility gradient downstream of FGF controls elongation of an amniote embryo. Nature 466, 248–252.

Blondel, V. D., Guillaume, J.-L., Lambiotte, R. and Lefebvre, E. (2008). Fast unfolding of communities in large networks. J. Stat. Mech. 2008, P10008.

Boyle-Anderson, E. A. T., Mao, Q. and Ho, R. K. (2022). Tbx5a and Tbx5b paralogues act in combination to control separate vectors of migration in the fin field of zebrafish. Dev Biol 481, 201–214.

Carver, E. A., Jiang, R., Lan, Y., Oram, K. F. and Gridley, T. (2001). The mouse snail gene encodes a key regulator of the epithelial-mesenchymal transition. Mol Cell Biol 21, 8184–8188.

Chacko, J., Ozadam, H. and Cenik, C. (2024). RiboGraph: an interactive visualization system for ribosome profiling data at read length resolution. Bioinformatics 40,.

Chaudhury, A., Cheema, S., Fachini, J. M., Kongchan, N., Lu, G., Simon, L. M., Wang, T., Mao, S., Rosen, D. G., Ittmann, M. M., et al. (2016). CELF1 is a central node in post-transcriptional regulatory programmes underlying EMT. Nat Commun 7, 13362.

Damon, B. J., Mezentseva, N. V., Kumaratilake, J. S., Forgacs, G. and Newman, S. A. (2008). Limb bud and flank mesoderm have distinct “physical phenotypes” that may contribute to limb budding. Dev Biol 321, 319–330.

Delgado, I., López-Delgado, A. C., Roselló-Díez, A., Giovinazzo, G., Cadenas, V., Fernández-de-Manuel, L., Sánchez-Cabo, F., Anderson, M. J., Lewandoski, M. and Torres, M. (2020). Proximo-distal positional information encoded by an Fgf-regulated gradient of homeodomain transcription factors in the vertebrate limb. Sci Adv 6, eaaz0742.

Delgado, I., Giovinazzo, G., Temiño, S., Gauthier, Y., Balsalobre, A., Drouin, J. and Torres, M. (2021). Control of mouse limb initiation and antero-posterior patterning by Meis transcription factors. Nat Commun 12, 3086.

Ebright, R. Y., Lee, S., Wittner, B. S., Niederhoffer, K. L., Nicholson, B. T., Bardia, A., Truesdell, S., Wiley, D. F., Wesley, B., Li, S., et al. (2020). Deregulation of ribosomal protein expression and translation promotes breast cancer metastasis. Science 367, 1468–1473.

ENCODE Project Consortium. (2012). An integrated encyclopedia of DNA elements in the human genome. Nature 489,.

Evdokimova, V., Tognon, C., Ng, T., Ruzanov, P., Melnyk, N., Fink, D., Sorokin, A., Ovchinnikov, L. P., Davicioni, E., Triche, T. J., et al. (2009). Translational activation of snail1 and other developmentally regulated transcription factors by YB-1 promotes an epithelial-mesenchymal transition. Cancer Cell 15, 402–415.

Fernández-Terán, M. A., Hinchliffe, J. R. and Ros, M. A. (2006). Birth and death of cells in limb development: a mapping study. Dev Dyn 235, 2521–2537.

Gros, J. and Tabin, C. J. (2014). Vertebrate limb bud formation is initiated by localized epithelial-to-mesenchymal transition. Science 343, 1253–1256.

Gros, J., Hu, J. K.-H., Vinegoni, C., Feruglio, P. F., Weissleder, R. and Tabin, C. J. (2010). WNT5A/JNK and FGF/MAPK pathways regulate the cellular events shaping the vertebrate limb bud. Curr Biol 20, 1993–2002.

Hagemann-Jensen, M., Ziegenhain, C., Chen, P., Ramsköld, D., Hendriks, G.-J., Larsson, A. J. M., Faridani, O. R. and Sandberg, R. (2020). Single-cell RNA counting at allele and isoform resolution using Smart-seq3. Nature Biotechnology 38, 708–714.

Haro, E., Delgado, I., Junco, M., Yamada, Y., Mansouri, A., Oberg, K. C. and Ros, M. A. (2014). Sp6 and Sp8 transcription factors control AER formation and dorsal-ventral patterning in limb development. PLoS Genet 10, e1004468.

Hasson, P., Del Buono, J. and Logan, M. P. O. (2007). Tbx5 is dispensable for forelimb outgrowth. Development 134, 85–92.

Heintzelman, K. F., Phillips, H. M. and Davis, G. S. (1978). Liquid-tissue behavior and differential cohesiveness during chick limb budding. J Embryol Exp Morphol 47, 1–15.

Hussey, G. S., Chaudhury, A., Dawson, A. E., Lindner, D. J., Knudsen, C. R., Wilce, M. C. J., Merrick, W. C. and Howe, P. H. (2011). Identification of an mRNP complex regulating tumorigenesis at the translational elongation step. Mol Cell 41, 419–431.

Infante, C. R., Park, S., Mihala, A. G., Kingsley, D. M. and Menke, D. B. (2013). Pitx1 broadly associates with limb enhancers and is enriched on hindlimb cis-regulatory elements. Developmental biology 374,.

Kania, A. and Jessell, T. M. (2003). Topographic motor projections in the limb imposed by LIM homeodomain protein regulation of ephrin-A:EphA interactions. Neuron 38, 581–596.

Kawakami, Y., Capdevila, J., Büscher, D., Itoh, T., Rodríguez Esteban, C. and Izpisúa Belmonte, J. C. (2001). WNT signals control FGF-dependent limb initiation and AER induction in the chick embryo. Cell 104, 891–900.

Korsunsky, I., Millard, N., Fan, J., Slowikowski, K., Zhang, F., Wei, K., Baglaenko, Y., Brenner, M., Loh, P.-R. and Raychaudhuri, S. (2019). Fast, sensitive and accurate integration of single-cell data with Harmony. Nature Methods 16, 1289–1296.

Lizarraga, G., Ferrari, D., Kalinowski, M., Ohuchi, H., Noji, S., Kosher, R. A. and Dealy, C. N. (1999). FGFR2 signaling in normal and limbless chick limb buds. Dev Genet 25, 331–338.

Loomis, C. A., Kimmel, R. A., Tong, C. X., Michaud, J. and Joyner, A. L. (1998). Analysis of the genetic pathway leading to formation of ectopic apical ectodermal ridges in mouse Engrailed-1 mutant limbs. Development 125, 1137–1148.

Mao, Q., Stinnett, H. K. and Ho, R. K. (2015). Asymmetric cell convergence-driven zebrafish fin bud initiation and pre-pattern requires Tbx5a control of a mesenchymal Fgf signal. Development 142, 4329–4339.

Martin, P. (1990). Tissue patterning in the developing mouse limb. Int J Dev Biol 34, 323–336.

McLean, C. Y., Bristor, D., Hiller, M., Clarke, S. L., Schaar, B. T., Lowe, C. B., Wenger, A. M. and Bejerano, G. (2010). GREAT improves functional interpretation of cis-regulatory regions. Nat Biotechnol 28, 495–501.

Min, H., Danilenko, D. M., Scully, S. A., Bolon, B., Ring, B. D., Tarpley, J. E., DeRose, M. and Simonet, W. S. (1998). Fgf-10 is required for both limb and lung development and exhibits striking functional similarity to Drosophila branchless. Genes Dev 12, 3156–3161.

Murray, S. A. and Gridley, T. (2006). Snail family genes are required for left-right asymmetry determination, but not neural crest formation, in mice. Proc Natl Acad Sci U S A 103, 10300–10304.

Musy, M., Flaherty, K., Raspopovic, J., Robert-Moreno, A., Richtsmeier, J. T. and Sharpe, J. (2018). A quantitative method for staging mouse embryos based on limb morphometry. Development 145,.

Newton, A. H. and Smith, C. A. (2024). Resolving the mechanisms underlying epithelial-to-mesenchymal transition of the lateral plate mesoderm. Genesis 62, e23531.

Newton, A. H., Williams, S. M., Major, A. T. and Smith, C. A. (2022). Cell lineage specification and signalling pathway use during development of the lateral plate mesoderm and forelimb mesenchyme. Development 149,.

Nishimoto, S., Wilde, S. M., Wood, S. and Logan, M. P. O. (2015). RA Acts in a Coherent Feed-Forward Mechanism with Tbx5 to Control Limb Bud Induction and Initiation. Cell Rep 12, 879–891.

Noguchi, K., Ishikawa, R., Kawaguchi, M., Miyoshi, K., Kawasaki, T., Hirata, T., Fukui, M., Kuratani, S., Tanaka, M. and Murakami, Y. (2017). Expression patterns of Sema3A in developing amniote limbs: With reference to the diversification of peripheral nerve innervation. Dev Growth Differ 59, 270–285.

Ohuchi, H., Nakagawa, T., Yamamoto, A., Araga, A., Ohata, T., Ishimaru, Y., Yoshioka, H., Kuwana, T., Nohno, T., Yamasaki, M., et al. (1997). The mesenchymal factor, FGF10, initiates and maintains the outgrowth of the chick limb bud through interaction with FGF8, an apical ectodermal factor. Development 124, 2235–2244.

Osterwalder, M., Speziale, D., Shoukry, M., Mohan, R., Ivanek, R., Kohler, M., Beisel, C., Wen, X., Scales, S. J., Christoffels, V. M., et al. (2014). HAND2 targets define a network of transcriptional regulators that compartmentalize the early limb bud mesenchyme. Dev Cell 31, 345–357.

Ozadam, H., Geng, M. and Cenik, C. (2020). RiboFlow, RiboR and RiboPy: an ecosystem for analyzing ribosome profiling data at read length resolution. Bioinformatics 36, 2929–2931.

Ozadam, H., Tonn, T., Han, C. M., Segura, A., Hoskins, I., Rao, S., Ghatpande, V., Tran, D., Catoe, D., Salit, M., et al. (2023). Single-cell quantification of ribosome occupancy in early mouse development. Nature 618, 1057–1064.

Poliak, S., Morales, D., Croteau, L.-P., Krawchuk, D., Palmesino, E., Morton, S., Cloutier, J.-F., Charron, F., Dalva, M. B., Ackerman, S. L., et al. (2015). Synergistic integration of Netrin and ephrin axon guidance signals by spinal motor neurons. Elife 4,.

Rallis, C., Bruneau, B. G., Del Buono, J., Seidman, C. E., Seidman, J. G., Nissim, S., Tabin, C. J. and Logan, M. P. O. (2003). Tbx5 is required for forelimb bud formation and continued outgrowth. Development 130, 2741–2751.

Robinson, M. D., McCarthy, D. J. and Smyth, G. K. (2009). edgeR: a Bioconductor package for differential expression analysis of digital gene expression data. Bioinformatics 26, 139.

Sedas Perez, S., McQueen, C., Stainton, H., Pickering, J., Chinnaiya, K., Saiz-Lopez, P., Placzek, M., Ros, M. A. and Towers, M. (2023). Fgf signalling triggers an intrinsic mesodermal timer that determines the duration of limb patterning. Nat Commun 14, 5841.

Sekine, K., Ohuchi, H., Fujiwara, M., Yamasaki, M., Yoshizawa, T., Sato, T., Yagishita, N., Matsui, D., Koga, Y., Itoh, N., et al. (1999). Fgf10 is essential for limb and lung formation. Nat Genet 21, 138–141.

Soshnikova, N., Zechner, D., Huelsken, J., Mishina, Y., Behringer, R. R., Taketo, M. M., Crenshaw, E. B., 3rd and Birchmeier, W. (2003). Genetic interaction between Wnt/beta-catenin and BMP receptor signaling during formation of the AER and the dorsal-ventral axis in the limb. Genes Dev 17, 1963–1968.

Sun, G., Lewis, L. E., Huang, X., Nguyen, Q., Price, C. and Huang, T. (2004). TBX5, a gene mutated in Holt–Oram syndrome, is regulated through a GC box and T-box binding elements (TBEs). Journal of Cellular Biochemistry 92, 189–199.

Suto, F., Ito, K., Uemura, M., Shimizu, M., Shinkawa, Y., Sanbo, M., Shinoda, T., Tsuboi, M., Takashima, S., Yagi, T., et al. (2005). Plexin-a4 mediates axon-repulsive activities of both secreted and transmembrane semaphorins and plays roles in nerve fiber guidance. J Neurosci 25, 3628–3637.

Tanigawa, Y., Dyer, E. S. and Bejerano, G. (2022). WhichTF is functionally important in your open chromatin data? PLoS Comput Biol 18, e1010378.

Tran, V., Papalexi, E., Schroeder, S., Kim, G., Sapre, A., Pangallo, J., Sova, A., Matulich, P., Kenyon, L., Sayar, Z., et al. (2022). High sensitivity single cell RNA sequencing with split pool barcoding. bioRxiv.

Urness, L. D., Bleyl, S. B., Wright, T. J., Moon, A. M. and Mansour, S. L. (2011). Redundant and dosage sensitive requirements for Fgf3 and Fgf10 in cardiovascular development. Dev Biol 356, 383–397.

Vaidya, A., Pniak, A., Lemke, G. and Brown, A. (2003). EphA3 null mutants do not demonstrate motor axon guidance defects. Mol Cell Biol 23, 8092–8098.

Vanlerberghe, C., Jourdain, A.-S., Ghoumid, J., Frenois, F., Mezel, A., Vaksmann, G., Lenne, B., Delobel, B., Porchet, N., Cormier-Daire, V., et al. (2019). Holt-Oram syndrome: clinical and molecular description of 78 patients with TBX5 variants. Eur J Hum Genet 27, 360–368.

Wada, N. (2011). Spatiotemporal changes in cell adhesiveness during vertebrate limb morphogenesis. Dev Dyn 240, 969–978.

Wanek, N., Muneoka, K., Holler-Dinsmore, G., Burton, R. and Bryant, S. V. (1989). A staging system for mouse limb development. J Exp Zool 249, 41–49.

Wurth, L., Papasaikas, P., Olmeda, D., Bley, N., Calvo, G. T., Guerrero, S., Cerezo-Wallis, D., Martinez-Useros, J., García-Fernández, M., Hüttelmaier, S., et al. (2016). UNR/CSDE1 Drives a Post-transcriptional Program to Promote Melanoma Invasion and Metastasis. Cancer Cell 30, 694–707.

Wyngaarden, L. A., Vogeli, K. M., Ciruna, B. G., Wells, M., Hadjantonakis, A.-K. and Hopyan, S. (2010). Oriented cell motility and division underlie early limb bud morphogenesis. Development 137, 2551–2558.

Xu, X., Weinstein, M., Li, C., Naski, M., Cohen, R. I., Ornitz, D. M., Leder, P. and Deng, C. (1998). Fibroblast growth factor receptor 2 (FGFR2)-mediated reciprocal regulation loop between FGF8 and FGF10 is essential for limb induction. Development 125, 753–765.

Ye, X., Tam, W. L., Shibue, T., Kaygusuz, Y., Reinhardt, F., Ng Eaton, E. and Weinberg, R. A. (2015). Distinct EMT programs control normal mammary stem cells and tumour-initiating cells. Nature 525, 256–260.

Zhu, M., Gu, B., Thomas, E. C., Huang, Y., Kim, Y.-K., Tao, H., Yung, T. M., Chen, X., Zhang, K., Woolaver, E. K., et al. (2024). A fibronectin gradient remodels mixed-phase mesoderm. Sci Adv 10, eadl6366.

